# Effects of organic and inorganic contaminants and their mixtures on metabolic health and gene expression in developmentally exposed zebrafish

**DOI:** 10.1101/2024.10.28.620642

**Authors:** Roxanne Bérubé, Matthew K. LeFauve, Aicha Khalaf, Darya Aminioroomi, Christopher D. Kassotis

## Abstract

Organic and inorganic chemicals co-occur in household dust, and these chemicals have been determined to have endocrine and metabolic disrupting effects. While there is increasing study of chemical mixtures, the effects of complex mixtures mimicking household dust and other environmental matrices have not been well studied and their potential metabolism disrupting effects are thus poorly understood. Previous research has demonstrated high potency adipogenic effects of residential household dust extracts using *in vitro* adipogenesis assays. More recent research simplified this to a mixture relevant to household dust and comprised of common co-occurring organic and inorganic contaminants, finding that these complex combinations often exhibited additive or even synergistic effects in cell models. This study aimed to translate our previous in vitro observation to an in vivo model, the developing zebrafish, to evaluate the metabolic effects of early exposure to organic and inorganic chemicals, individually and in mixtures. Zebrafish embryos were exposed from 1 day post fertilization (dpf) to 6 dpf, then metabolic energy expenditure, swimming behavior and gene expression were measured. Globally, we observed that most mixtures did not reflect the effects of individual chemicals; the BFR mixture produced a less potent effect when compared to the individual chemicals, while the PFAS and the inorganic mixtures seemed to have a more potent effect than the individual chemicals. Finally, the environmental mixture, mimicking household dust proportions, was less potent than the inorganic chemical mix alone. Additional work is necessary to better understand the mixture effect of inorganic and organic chemicals combined.

## 1. Introduction

Obesity is rising in incidence worldwide and is a global health issue among adults and children. Obese or overweight individuals have increased risks of type 2 diabetes, cardiovascular disease, dyslipidemia and metabolic syndrome. Recent studies demonstrated that energy balance and genetics are not the only factors explaining the global increasing obesity incidence (Egusquiza & Blumberg, 2020; Lustig et al., 2022). Environmental factors, such as stress and gut microbiome composition, are known to affect obesity, and more recently endocrine disrupting chemicals (EDCs) have been suspected to affect weight gain. EDCs disturb molecular mechanisms and pathways associated with weight gain, adiposity, glucose and insulin signaling (Heindel et al., 2017).

As early as gestation, but through the lifespan, humans are chronically exposed to various chemicals, from our diet and our environment (Dallaire et al., 2003; Houlihan et al., 2005). One important chronic source of exposure is household dust, which can be ingested or inhaled, and contains thousands of organic and inorganic chemicals from consumer products, cookware and building materials, which co-occur in complex mixtures at high frequency (Hammel et al., 2019; Kassotis et al., 2021; Phillips et al., 2018). These chemicals include poly and -perfluoroalkyl substances (PFAS), brominated flame retardant (BRFs), polychlorinated biphenyls (PCBs), and metals (lead, cadmium, arsenic, etc.), among others. PFAS are persistent synthetic compounds that are ubiquitously measured in various human tissues, such as blood, serum, breast milk, etc. (Kannan et al., 2004; LaKind et al., 2023). An increasing number of studies have demonstrated the adverse effects of PFAS on human health: epidemiological studies have found relation with PFAS exposures and kidney and testicular cancers (Barry et al., 2013), thyroid disease (Melzer et al., 2010), and adiposity (Timmermann et al., 2014). They act as endocrine disrupting chemicals by leading to reproductive and developmental toxicity (González-Alvarez et al., 2024) and have been associated with metabolic diseases (Momo et al., 2024).

BFRs are persistent organic pollutants (POPs) used to inhibit fire that are frequently found in furniture, electronic components and firefighting foam (D’Hollander et al., 2010). They are particularly relevant for the Michigan population after an agricultural accident in 1973-1974 led to a contamination of food supply and higher levels of BFRs, particularly PBB-153, are still present in exposed residents four decades later (González-Alvarez et al., 2024; Hoffman et al., 2023). Studies from the Michigan cohort demonstrated that exposures to BFRs led to perturbations in the metabolic pathways, increased inflammation and oxidative stress. Other studies have demonstrated that BFR exposures induced testicular toxicity in mice (Zhang et al., 2022) and affected lipid metabolism and glucose through PPARs signaling and the mTOR pathway in HepG2 cells (Casella et al., 2022).

Human activity contributes to the release of inorganic contaminants such as lead, cadmium, and arsenic into the environment, particularly through our diets and our homes. Lead exposures are primarily from ingestion and inhalation of contaminated substances and affect neurodevelopment, behavior and cognitive functions (Al Osman et al., 2019). The mechanisms of lead toxicity come from its ability to inhibit key enzymes in the heme synthesis pathway and antioxidant enzymes (Flora et al., 2012). Lead can substitute calcium ions (Ca^2+^), accumulate in bones (Barbosa et al., 2005), cross the blood brain barrier (Bradbury & Deane, 1993) and affects neurotransmitter release (Bressler et al., 1999). Cadmium exposures also arise from ingestion or inhalation of contaminated sources, such as household dust, food, and water (Genchi et al., 2020). Cadmium accumulates mainly in the kidneys, liver, and intestines (Satarug et al., 2023), where it creates organ dysfunctions and diseases (Sabolić et al., 2010). The toxicity of cadmium is due to its capacity to disrupt mitochondrial proteins, inhibit the electron-transfer chain, and can induce DNA damage and disrupt DNA methylation and repair (Pizzino et al., 2014). Lastly, arsenic is used in a vast array of consumer products, in agriculture and in medical treatments, despite its well-described toxicity (Paul et al., 2023; Tchounwou et al., 2004). Similarly to cadmium and lead, arsenic acts mainly through induction of oxidative stress, increasing oxidative damage to lipid and DNA, can lead to acute or chronic toxicity at the organ or tissue level (e.g., neurologic, cardiovascular, respiratory, etc.) (Fatoki & Badmus, 2022) and can induce cancer of the skin, bladder, kidney or lung (Rahaman et al., 2021).

Overall, the main effects of these substances individually are well-understood, and they have been reported as endocrine and metabolic disrupting chemicals. Despite individual contaminant reports, very few studies have considered the effects of organic and inorganic chemical mixtures on metabolic health, which is more representative of environmental exposures (Wattigney et al., 2022). Better understanding realistic mixtures of contaminants is important as growing research has documented that chemicals can act in concert to elicit greater effects than could be predicted based on individual component chemical effects alone (Martin et al., 2021; Rajapakse et al., 2002; Silva et al., 2002). Due to the presence of these combined chemicals in household dust, the endocrine disrupting effects of dust exposures requires further investigation. We recently demonstrated the effects of organic and inorganic chemicals on metabolic processes and pathways, by measuring receptor activity, adipocyte differentiation and lipid accumulation *in vitro* (Bérubé et al., 2023). PFAS, BFRs, and inorganics all promoted adipogenesis in human mesenchymal stem cells and the effects of combinations of these (class-based mixtures, organic + inorganic mixtures) all produced significantly greater effects than would be expected based on individual component chemicals. These interactions were often deemed putatively synergistic based on mixture analysis modeling. These apparent greater than anticipated effects across diverse chemical classes suggested that these combinations could potentially account for the robust adipogenic activity exhibited by small concentrations of residential household dust samples reported previously (Kassotis et al., 2021; Kassotis et al., 2017; Kassotis et al., 2019). In particular, combinations of PFAS, BFRs, and organics/inorganics interacted to promote greater *in vitro* adipogenic effects.

To follow up on our *in vitro* mixture assessment, this current project aimed to observe metabolic effects of those same chemicals and mixtures *in vivo* using a vertebrate model, the zebrafish (*Danio rerio*), to determine translation to whole organismal metabolic health. We aimed to measure toxicity, metabolic activity, swimming behavior and expression of genes involved in lipid and glucose metabolism, detoxification process, and cellular receptors in 6-day post fertilization (dpf) zebrafish larvae developmentally exposed to organics, inorganics, and their mixtures.

## 2. Materials and Methods

### 2.1. Chemicals

Chemicals used are described in detail in Table 1. Stock solutions were prepared in 100% DMSO (Sigma cat # D2650) and stored at -20 °C between uses. The mixtures were prepared with an equimolar concentration of each chemical and the environmentally relevant mixture were prepared with the inorganic 100-fold higher than the organic (e.g., 10 µM environmental mix = 10 µM of each inorganic contaminant and 100 nM of each organic contaminant), modeling concentrations previously reported in household dust samples (Bérubé et al., 2023; Kassotis et al., 2021).

**Table 1.** Organic and Inorganic Constituent Contaminants.

| CONTAMINANT | ACRONYM | CAS # | SUPPLIER | CATALOG # |
| --- | --- | --- | --- | --- |
| Perfluorooctanoic acid | PFOA | 335-67-1 | Sigma | 33824-100MG |
| Perfluorooctanesulfonic acid | PFOS | 1763-23-1 | SCBT | sc-235283B |
| 2,2',4,4'-tetrabromodiphenyl ether | BDE-47 | 5436-43-1 | Sigma | 91834-10MG |
| 2,2',4,4',5,5'-hexabromobiphenyl | PBB-153 | 59080-40-9 | Accustandard | B-153N-5mg |
| Lead acetate | Pb | 6080-56-4 | Sigma | 316512-5G |
| Sodium Arsenite | As | 7784-46-5 | Sigma | S7400-100G |
| Cadmium chloride | Cd | 233-296-7 | Sigma | 202908-10G |

**Table 2.**
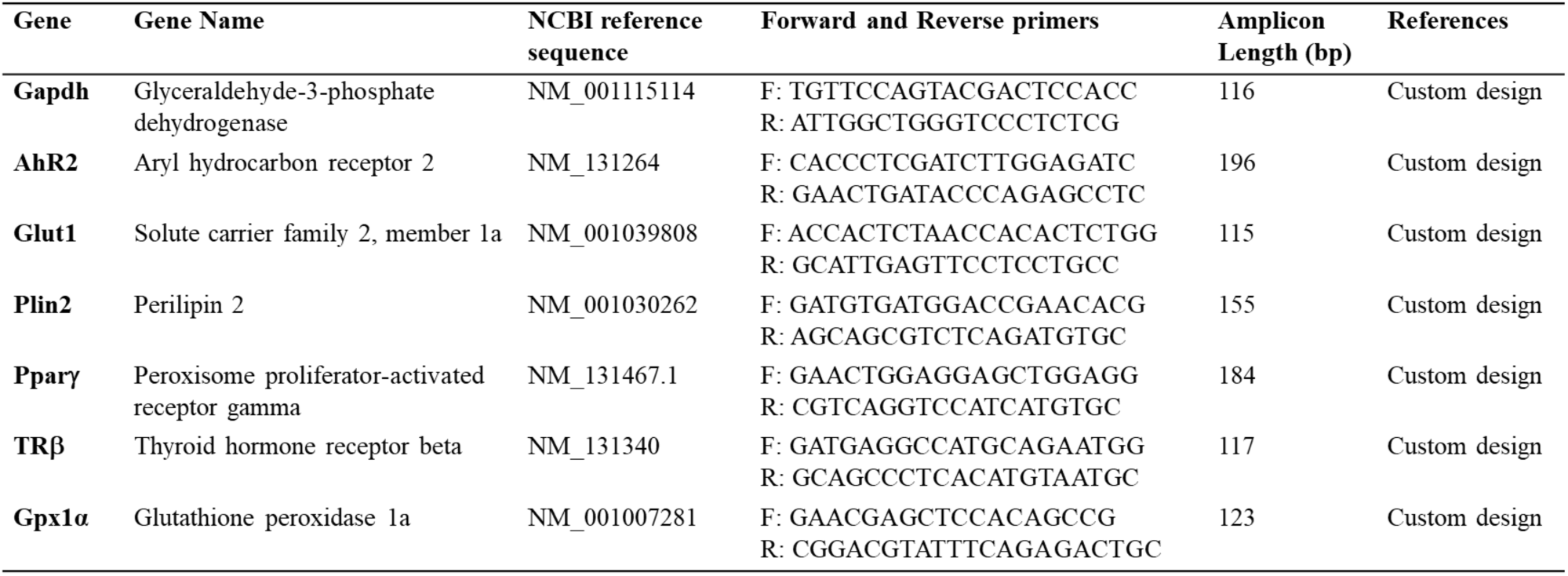
Genes of interest and specific primer parameters.

### 2.2. Zebrafish housing and care

Wildtype AB zebrafish (*Danio rerio*) were housed and cared for according to standard protocols (Westerfield 2000). and best ethical practices as approved by the Wayne State University Institutional Animal Care and Use Committees (protocol # IACUC-20-06-2408). To generate embryos, AB adult zebrafish were paired in breeding chambers, separating males and females overnight. Water was changed and separators were removed at time of lights on, and embryos were collected at the conclusion of the spawning event. Embryos were cleaned and stored overnight in embryo media (EM) with methylene blue (0.1%). Zebrafish were fed from 6 dpf with GEMMA Micro 75 (Skretting, USA) twice per day until 15 dpf and maintained in crystallizing jars in 20-30 mL of EM with media changes at least every other day. At 15 dpf, fish were transferred to a flow-through system in 4.5L tanks, and they were switched to GEMMA Micro 150 until 30 dpf.

### 2.3. Zebrafish exposures

Prior to the exposures, a toxicity test was performed with each chemical to determine sublethal concentration. For all chemicals the concentrations inducing minimal mortality (>10%) were 100, 10, 1 and 0.1 nM. At approximately 24 hours (1 dpf) following spawns, viable embryos were separated out into 500 mL glass jars in 30-50 mL of EM for chemical exposures (n=30-50 individual embryos per chemical test concentration). Chemical exposures were performed in EM using individual chemical stocks at 0.1% dimethylsulfoxide (DMSO) vehicle. Zebrafish were exposed from 1 dpf through 6 dpf, with media and test chemical changes made every 24 hours to ensure consistent dosing. Concentrations were not determined in the dosing medium; as such, they should be considered as nominal concentrations only. As of 6 dpf, exposure media was replaced with fresh EM without test chemicals. Fish were subsequently aged out to 30 dpf to perform additional analyses, at which time they were sacrificed and snap frozen.

### 2.4. Energy expenditure measurements

Energy expenditure was measured using the alamar blue assay, adapted from previously published protocols (Reid et al., 2018; Renquist et al., 2013). Briefly, following chemical exposures, 6 dpf zebrafish larvae were transferred to fresh EM without added chemicals or methylene blue. Three larvae per treatment and control group were transferred in one well of a 24-well black clear-bottom microtiter plate (n=3 well per group, and four separate exposure experiments). EM was removed from wells and replaced with 1 mL of alamar blue solution (0.2X alamarBlue™ Cell Viability Reagent in filtered EM). Plates were read immediately using an iD5 Molecular Devices plate reader using 530/590 excitation/emission wavelengths and a second read was obtained approximately 16 hours later. Between reading, plates were kept in a 28°C incubator, in the dark. Metabolic activity was determined by the difference in fluorescence units (16-hour read – initial read) normalized to the difference in fluorescence from the DMSO control group.

### 2.5. Behavior assessment

Larval activity, as assessed by swim distance in light and dark cycles, was automatically quantified using Noldus Ethovision (version XT 16; (Noldus et al., 2001)) during a 45-minute test period. Briefly, six larvae from each control and exposure group were placed individually in a 24-well plate, in fresh EM and in the absence of test chemicals and were allowed to acclimate to a sound-insulated, temperature-controlled (26°C), and light controlled testing chamber. All larvae were subjected to a 5-minute acclimation, followed by two cycles consisting of a 10-minute period of light and then a 10-minute period of dark (Fitzgerald et al., 2021; Stewart et al., 2011). Movement of 24 individual larvae was measured using an auto-detect feature of Ethovision, with all movement data binned into 60-second intervals. The resulting data were verified using Ethovision before statistical analysis and any movement >20 cm per minute was removed, as they were determined to be software artifacts by visual observations of the tracking video files. Raw data were exported, and average total distance moved (cm/min) was analyzed in Graphpad Prism 9.0. Statistical differences were calculated using a non-parametric Kruskal-Wallis, followed by a Dunn’s uncorrected multiple comparison test. Each chemical in each light/dark period was tested against the DMSO control group. The assay was performed after three different exposures experiments, for a total of 18 larvae per treatment (n=6 fish per group, and 3 separate exposure experiments).

### 2.6. Gene expression

At the end of the exposure, pools of 10 larvae in each treatment group were snap-frozen with liquid nitrogen (n=5 pools of 10 larvae per treatment each). RNA was isolated with the Qiagen miRNeasy Micro kit by following the manufacturer’s protocol. Briefly, Qiazol® lysis reagent and two stainless steel beads were added to the samples for homogenization with the Bullet blender (3 cycles of 30 sec at speed 4). The samples were incubated for 5 min at room temperature, chloroform was added, and the mixture was vortexed and incubated an additional 2 min. The samples were centrifuged (15 min, 12 000 g, 4 °C) and the aqueous phase was collected in a new tube containing 100% ethanol. The samples were transferred to a spin column in a collection tube. Washing steps with the recommended buffers were performed and finally RNA was eluted twice with RNase-free water. The RNA was quantified with a Nanodrop (ThermoFisher) and diluted for the following steps. The gDNA was removed and cDNA was synthesized, using the iScript^TM^ gDNA Clear cDNA Synthesis kit (Bio-Rad). Gene expression was measured using the SsoAdvanced Universal SYBR® Green Supermix (Bio-Rad). To ensure accuracy, samples without reverse-transcriptase and no-template controls were included. The relative expression of each gene (*ahr*, *glut1*, *plin2*, *pparγ*, *trβ,* and *gpx1α*) was normalized using the expression of *gapdh* as the reference gene and calculated with the 2^-ΔΔCT^ method, using the Ct mean of the reference gene. Primer sequences and information are listed in Table S2. All primers were tested with a standard curve to ensure efficiency between 90 and 100% and a R^2^ of at least 0.98.

### 2.7. Statistical Analysis

Data are presented as means ± SEM from 3 to 6 technical replicates (individual larvae or pool) from three or four independent biological replicates (independent spawning events and exposure). Two-way Kruskal-Wallis with Dunn’s uncorrected multiple comparisons test was performed to determine significant differences across concentrations and relative to DMSO-control fish (p<0.05 considered significant). Statistical comparisons and figures were made using GraphPad Prism 9.0.

## 3. Results

### 3.1. Toxicity

Toxicity of 6 dpf larvae was measured as percentage of survival (Fig. 1). No significant difference in survival was measured, and 95 to 100% survival was observed for all treatments. Preliminary toxicity testing demonstrated that higher concentrations (10 and 1 uM) of each individual chemical induced significant mortality in most treatments, from 50% to 100%, and they were removed from this study.

**Figure 1:**
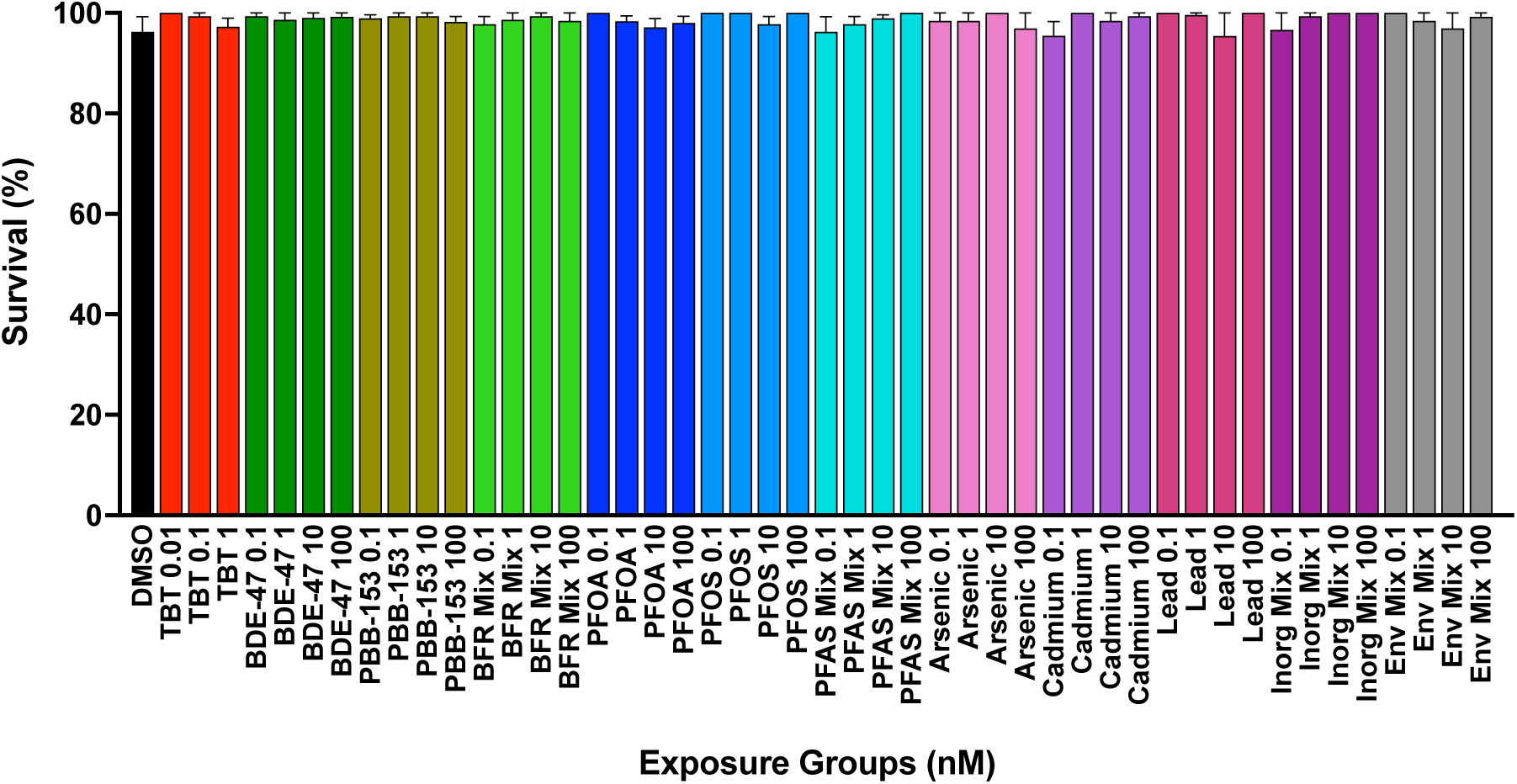
Zebrafish larvae survival (%) at 6 dpf after exposures to test chemicals and mixtures. Zebrafish were developmentally exposed to control chemicals, each individual organic and inorganic chemical, and their mixtures. Immediately following exposure, at six days post fertilization, survival was calculated (percent surviving at 6 dpf) and significant differences were calculated by comparing the survival of treated fish to DMSO treated fish using Kruskal-Wallis and multiple comparisons tests. Data are presented as mean ± SEM.

### 3.2. Energy expenditure measurements

Energy expenditure of zebrafish larvae exposed to organic and inorganic chemicals and their mixtures was measured with the alamar blue assay and is presented in Fig. 2 as a relative change in fluorescence. A significant increase in metabolic activity in comparison to the DMSO control group was observed in zebrafish larvae exposed to PBB-153 at 10 and 100 nM and tended to increase in the BFR mixture at 100 nM (p < 0.1). In contrast, significant decreases in metabolic activity were induced by exposures to PFOS and cadmium at 0.1 nM, and by the PFAS mixture and the environmental mixture at 100 nM. The metabolic activity tended to decrease in fish exposed to PFOA at 100 nM. All other treatments, including the positive control TBT, did not induce any significant change in metabolic activity.

**Figure 2:**
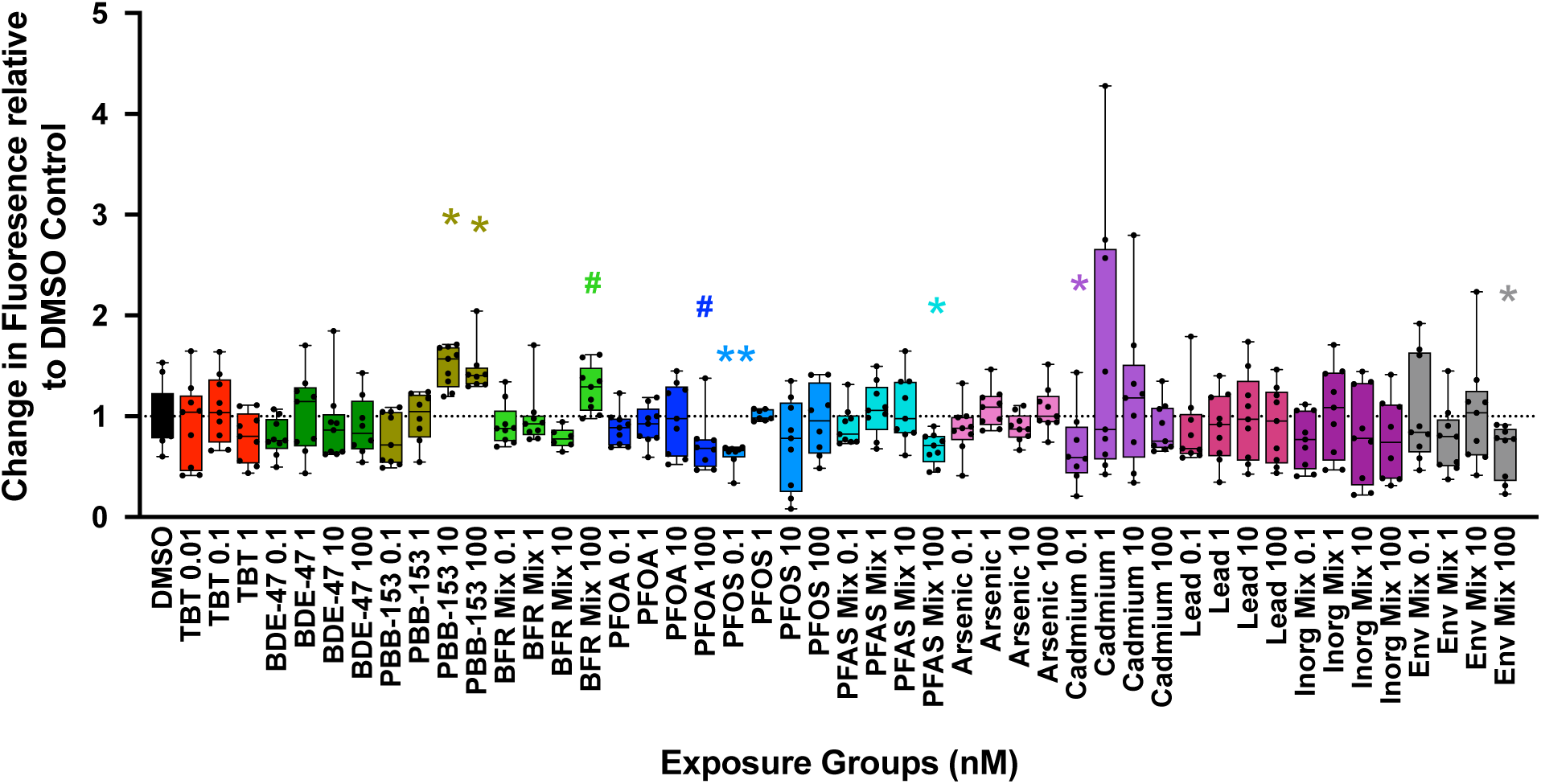
Metabolic activity in zebrafish developmentally exposed to organic and inorganic chemicals and their mixtures. Zebrafish were developmentally exposed to control chemicals, each individual organic and inorganic chemical, and their mixtures. Immediately following exposure, at six days post fertilization, metabolic activity was measured using the alamar blue assay. Significant differences were calculated by comparing treated fish responses with DMSO treated fish. Data are presented as mean and quartiles. N = 3 wells of 3 fish from 3 independent breeding event and exposure. *p < 0.05, **p < 0.01 as per Kruskal–Wallis test with Dunn’s multiple comparisons.

### 3.3. Behavior assessment

Distance traveled by zebrafish larvae exposed to organic, inorganic contaminants, and their mixture was measured in a 45-minute light/dark assay (Fig. 3). Hyperactivity in both dark periods was observed for each BFR individually (BDE-47 at 1 and 100 nM; PBB-153 at 1, 10 and 100 nM), however, the BFR mixture did not induce hyperactivity (Fig. 3A-C). The PFAS mixture induced hyperactivity in both dark periods at 0.1, 10, and 100 nM (Fig. 3F), while for single PFAS only PFOA at 0.1 nM (Fig. 3D) induced hyperactivity in both dark periods (PFOA 0.1 nM: Fig. 3D-E). However, PFOA at 100 nM, PFOA at 1 nM, and PFOS at 10 nM (Fig. 3E) increased swimming distance in exposed larvae in only one dark period. The inorganic chemicals, specifically cadmium at 0.1 and 10 nM (Fig. 3H) induced hyperactivity in both light and dark periods of the assay, which was also reflected in the inorganic mixture at 10 nM and for the first cycle only at 100 nM. The arsenic exposures induced hyperactivity for only one concentration (0.1 nM) and in one dark period, while lead exposures induced hyperactivity in both dark periods, except at 100 nM, which induced hyperactivity in one dark period only.

**Figure 3:**
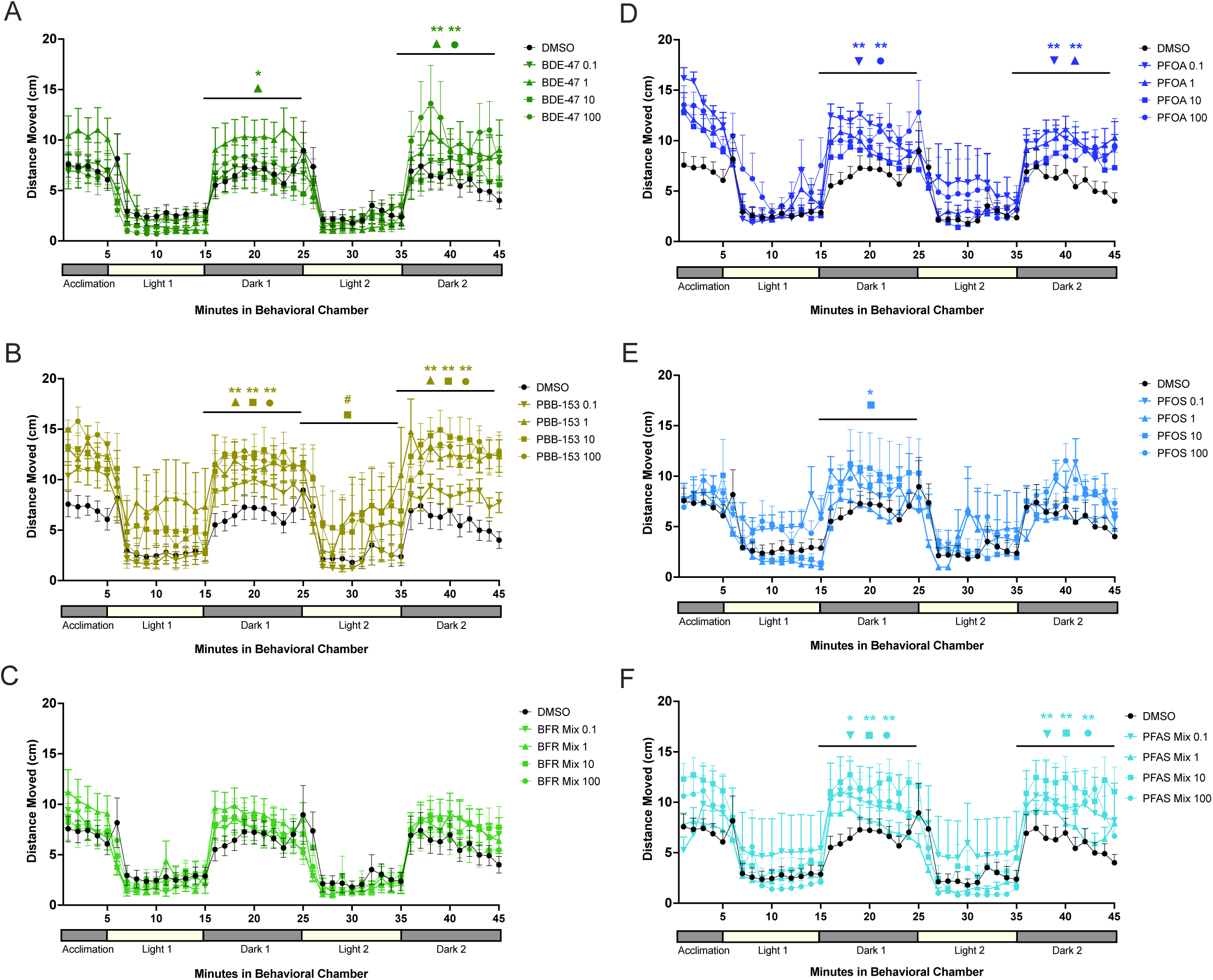
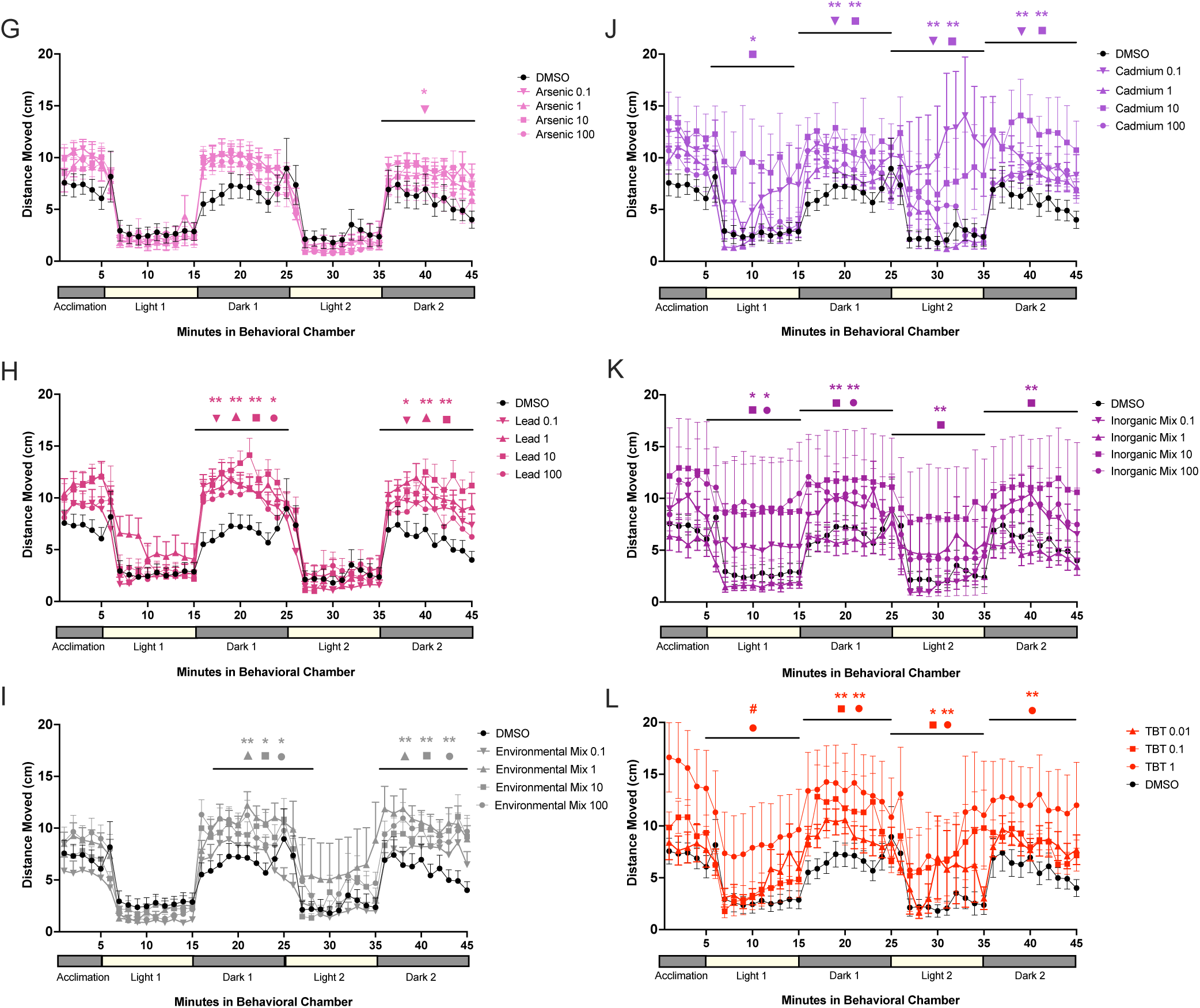
Total distance traveled during light/dark neurodevelopmental testing. Zebrafish were developmentally exposed to control chemicals, each individual organic and inorganic chemical, and their mixtures. Immediately following exposure (6 dpf), activity was tracked using Noldus Ethovision software. Six fish per treatment were transferred individually into wells of a 24-well plate. The total activity was tracked using a 5 min acclimation period followed by two cycles of one ten-minute light and one ten-minute dark period. Significant differences were calculated by comparing the total swimming distance in each ten-minute period for each chemical to DMSO treated fish. N = 6 individual fish from 5 independent breeding events and exposure. *p < 0.05, **p < 0.01 as per Kruskal–Wallis test with Dunn’s multiple comparisons.

### 3.4. Gene expression

We measured the expression of genes in metabolic health signaling (*pparγ*: regulates fatty acid storage and glucose metabolism; *glut1*: facilitates glucose transport; and *plin2*: adipose differentiation-related protein), hormone receptor (*trβ*: thyroid hormone receptor); and in detoxification metabolism (*ahr2*: transcription factor in various signaling processes; and *gpx1α*: detoxification/antioxidant enzyme) (Fig. 4). Fish exposed to TBT at 1 nM had decreased expression of most genes (*pparγ*, *plin2*, *ahr2*, and *gpx1α*). No significant change in *pparγ* expression was induced by our exposures, however PBB-153 at 100 nM tended to cause a reduction in *pparγ* expression (p < 0.1, Fig. 4A). Exposure to cadmium at 100 nM induced a significant increase in *glut1* expression, while it tended to increase following exposure to BDE-47 at 1 nM, PFAS mix at 10 nM and lead at 100 nM (Fig 4B). The expression of *plin2* tended to decrease when fish were exposed to PBB-153 at 100 nM and cadmium at 1 nM (Fig. 4C). Next, the expression of *trβ* was significantly increased by the inorganic mixture at 1 nM and tended to increase with the exposures to PFOA at 1 nM, PFOS at 100 nM, PFAS mix at 10 and 100 nM, and the inorganic mix at 10 nM (Fig. 4D). Exposure to PBB-153 at 10 nM induced a decrease in *ahr2*, while it tended to decrease after exposure to 100 nM of PBB-153 (Fig. 4E). Finally, the expression of *gpx1α* was significantly increased by arsenic at 1 nM, tended to increase with the inorganic mix at 1 nM, and tended to decrease with exposures to BDE-47 at 100 nM and PBB-153 at 10 and 100 nM (Fig. 4F). No gene expression was affected after exposure to any concentration of the environmental mixture.

**Figure 4.**
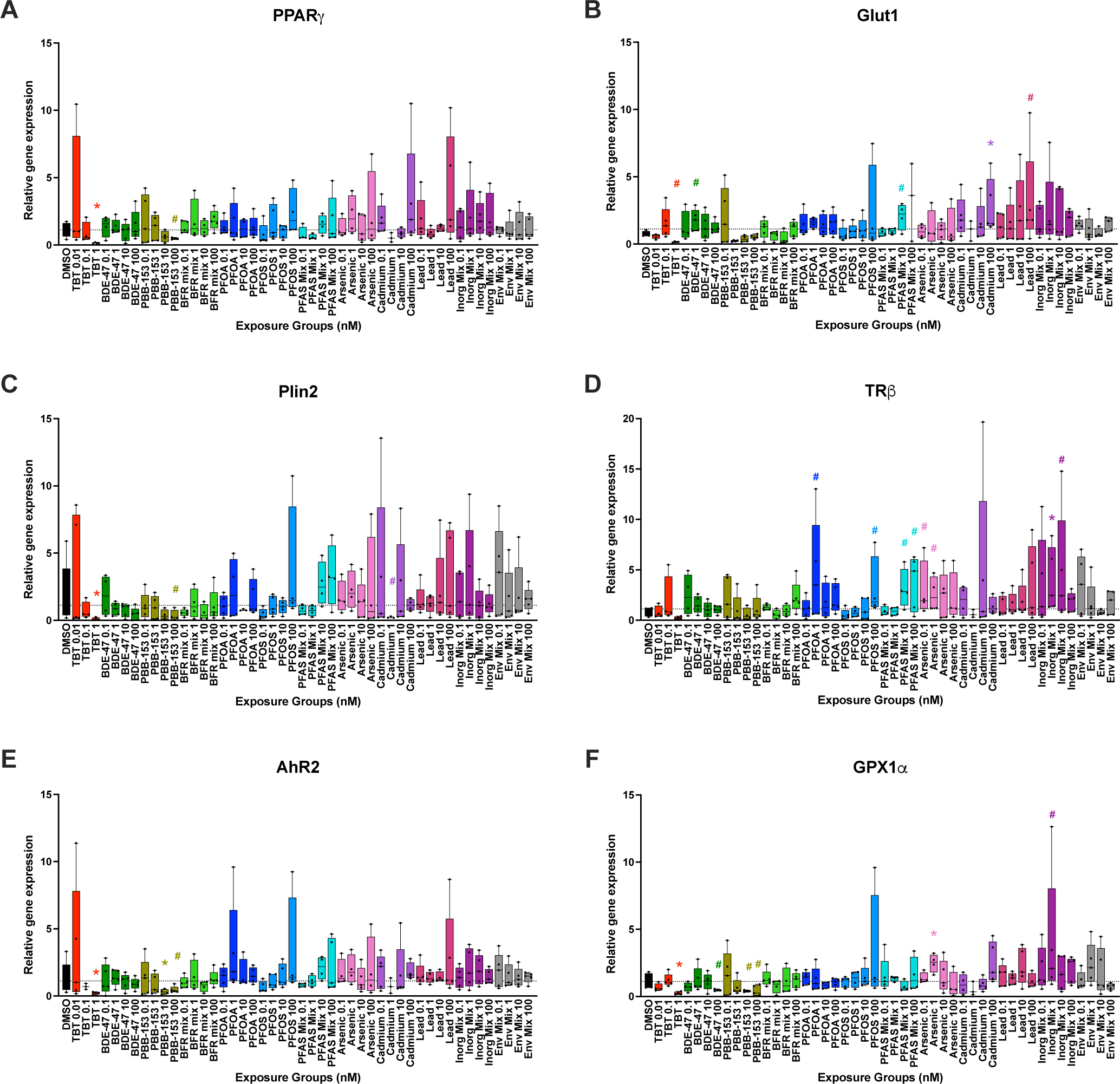
Gene expression in zebrafish developmentally exposed to organic and inorganic chemicals and their mixtures. Zebrafish were developmentally exposed to control chemicals, individual organic and inorganic chemicals, and to mixtures of PFAS, BFRs, inorganics and an environmental mixture. Relative gene expression was calculated using the 2^-ΔΔCT^ method after qPCR analysis. For each gene, the relative expression in the treated fish was compared to the relative expression in DMSO treated fish. *p < 0.05, as per Kruskal–Wallis test with Dunn’s multiple comparisons. # indicates 0.05 < p < 0.1 N = 3 to 5 pools of 10 fish from independent breeding event and exposure.

## 4. Discussion

In this study, we evaluated the toxicity and effects on metabolic health of 6 dpf zebrafish larvae developmentally exposed to organic and inorganic chemicals, individually and in mixtures, representing realistic human co-exposures reflective of those encountered in residential house dust. The effects of the mixtures in zebrafish were complex and did not always reflect expected combined effects on early-life developmental endpoints; in short, often individual chemicals had more significant effects than mixtures. This was not universally true, with some mixtures exhibiting greater effects on specific endpoints than individual component chemicals, and further research is needed to better elucidate why some combinations and not others act cooperatively towards these metabolic health endpoints and others do not (or may even act antagonistically in combination).

We evaluated the effect of two brominated flame retardants (BFRs), BDE-47 and PBB-153, individually and in an equimolar mixture. Our results demonstrated that BDE-47 induced a moderate increase in swimming activity in the dark, although it did not affect the metabolic activity. PBB-153 induced an increased metabolic activity at high concentrations (10 and 100 nM) and increased the swimming activity of exposed fish in the dark for the three highest concentrations. The increase in metabolic activity induced by PBB-153 was associated with an increase in swimming behavior. However, for BDE-47, the changes in the swimming activity were not mirrored by the metabolic activity assessment. The swimming activity was only observed consistently for one concentration of BDE-47, and it is possible that the modifications in activity over a small time period may be too low and cannot be observed in the metabolic activity over 16 hours. A previous study has demonstrated that BDE-47 has a low ability to cross the chorion and accumulates in the embryo rapidly after hatching (Liu et al., 2015). This may explain its lower efficacy in most behavioral endpoints compared to PBB-153.

For both assays, the BFR mixture did not completely reflect the individual chemical effects: the high mixture concentration tended to affect the metabolic activity and did not affect the swimming activity. In a similar study, zebrafish embryos were exposed to BDE-47 (Chackal et al., 2022), among other treatments, and their study measured the swimming behavior and the metabolic activity. They used the alamar assay combined with a high-precision respirometry assay to measure the oxygen consumption rate. This study demonstrated that the oxygen consumption rate did not relate with the Alamar assay measurements (measured similarly to our study), with the BDE-47 exposure inducing a higher oxygen consumption rate compared to the control, but no difference observed in the alamar assay. Furthermore, Chackal et al. (2022) did not find difference between the BDE-47 exposed group and the control group, in agreement with our data, though did find a difference in swimming speed. Their results compared to ours seems to demonstrate that the alamar assay may not be the most precise to evaluate metabolic activity and that BDE-47 induces moderate behavioral modification. There are multiple studies on the effects of PBB-153 exposure in the Michigan population after an accidental contamination of the food chain. These studies demonstrated that PBB-153 bioaccumulates (Chang et al., 2020) and can lead to epigenetic alterations affecting the next generation in humans (Curtis et al., 2020; Greeson et al., 2020). *In vitro* assays in trout hepatocytes demonstrated that exposures to BDE-47 and PBB-153 caused an increase in vitellogenin and then a sharp decrease at higher concentrations. A similar response was observed with the EROD assay, which measures the detoxifying capacity of the cells (Nakari & Pessala, 2005). Taken together, these results demonstrated that individual BFRs can be neurotoxic, affect development, and impact behavior and metabolism. However, their combined toxicity is not well understood, and more research is necessary as the mixtures appeared to act antagonistically towards the endpoints measured here (though this requires substantiation with broader dose response assessments and formal mixture effect calculations).

Metabolic activity was not modified by most PFAS exposures, with decreases noted for only PFOS at 0.1 nM and for the PFAS mix at 100 nM. In contrast, swimming activity in the dark was increased by the PFAS mixture. Effects on swimming activity appeared to be additive, as some increased activity was induced by the individual chemicals but a consistent increase in the dark was induced by the three highest PFAS mixture concentrations. Lastly, *TRβ* and *glut1* expression tended to increase with PFAS exposures. Previous studies, as reviewed in (Cao & Ng, 2021), have demonstrated that PFAS, including PFOA and PFOS can accumulate in the brain of mammals and various fish species, with longer chain PFAS accumulating at higher concentrations. PFOS contains a sulfonate group, which can form more hydrogens bonds with amino acids and increases its accumulation in tissues compared to PFOA, which contains a carboxylate group (Wen et al., 2019). Our results demonstrated that PFOA induced slightly greater effects than PFOS in the behavior testing, although the mixture was the most potent in affecting swimming behavior. The neurotoxicity of PFAS can be explained by their ability to affect calcium homeostasis in neurons and induce neuronal excitement and/or neuron injury (Liao et al., 2008; Liu et al., 2011) and/or the thyroid receptor beta antagonism we previously reported for these PFAS *in vitro* (Bérubé et al., 2023). Globally, PFOA and PFOS induced individual effects, but their combined mixture induced more potent effects, suggesting mixture additivity or potential synergism, which will require additional research with broader dose responses to comprehensively model and predict deviations from expected mixture effects.

For the inorganic contaminants, our results demonstrated that metabolic activity was decreased by cadmium at 0.1 nM. We also observed that the swimming activity was increased in the dark by lead, and in both light and dark periods by cadmium. The increased swimming activity in both periods was also present in fish exposed to the mixture of inorganics. Lastly, the gene expression of *glut1* was modified by the highest cadmium concentration, tended to increase with high concentrations of lead, but was not modified by any other inorganic chemicals or the mixture. The inorganic mixture did promote changes in expression of *TRβ* and *GPX1α*. Developmental lead exposures were previously demonstrated to cause neurological damage (Liu et al., 2023) and these effects occurred mostly before the complete formation of the blood-brain barrier, which happens between 3 and 5 dpf in zebrafish (Jeong et al., 2008; O’Brown et al., 2019), which is during the exposures conducted here. Our results demonstrated that inorganic chemicals affected swimming behavior, potentially via neurotoxicity. Our results are corroborated by other work, where lead-exposed zebrafish had an increased swimming speed and memory and learning deficits were observed at later life stages (Chen et al., 2012). Among the three inorganic chemicals evaluated in this study, cadmium induced the greatest effect, with cadmium alone inducing hyperactivity in both the dark and light periods. Arsenic did not cause any notable effect in any endpoints, which may be due to its inability to cross the chorion (Olivares et al., 2016), while cadmium and lead have been shown to cross the chorion and affect its structure and protection capacity (Cheng et al., 2000; Zhang et al., 2011). Lastly, the mixture seemed to be more potent than the individual chemicals, but not for all the measured endpoints, suggesting complex effects of the inorganics on the measured metabolic health outcomes.

Lastly, the environmental mixture (containing 100-fold higher concentrations of inorganics compared to the organics) demonstrated decreased metabolic activity at the high concentration and increased swimming activity across most concentrations. However, the increase in swimming activity in the light period, induced by cadmium individually, was not reflected in the mixture exposures. This suggests potential mitigation of the inorganic effects by the organic constituents, even though they were present at considerably lower concentrations. Considering their molecular charges, there is high probability that these chemicals interact and bind in water (Wang et al., 2023; Xing et al., 2022). Both lead acetate and cadmium chloride, the two most potent metals used in this study, dissociate in water to form lead ions (Pb^2+^) and cadmium ions (Cd²⁺), while PFOA and PFOS have negative functional groups (-COO^-^ and -SO3^-^, respectively). If these compounds bind together, their toxicity could be affected and could possibly decrease interactions with cellular components, decreasing the immediate toxicity. One study demonstrated that these interactions can affect the transport of these chemicals and their bioavailability in the environment depending on the solution chemistry and the presence of dissolved organic matter (Wang et al., 2023), However, this needs more research to understand fully the mixture effects of these compounds and how their chemical interactions affect their toxicity.

The gene expression measured in this study generally lead to few significant changes from controls. This could be explained by the tissue-specific expression of most genes evaluated in this study. We used pools of whole embryos, which may have caused a dilution of the localized and precise changes in expression. For example, previous work has demonstrated the localization of the thyroid hormone receptor genes in developing embryos and demonstrated that it is tissue-specific, varies widely depending on the developmental stage, and stabilizes around 48 to 72 hpf (Marelli et al., 2016). Gene expression at later timepoints, particularly in specific organs, or potentially even single-cell RNA-seq at these early timepoints may be able to discriminate more effects than we observed here.

Furthermore, contrary to our expectations and previous literature (Zhou et al., 2023) obtained responses between the metabolic and behavior assays were not often related. The swimming behavior assay was a 45-minute test period, while the Alamar blue assay reflects the average metabolic activity over 16 hours, completely in the dark, which may explain some of the difference between both assays; fish in the same condition over a long period, either light or dark, generally decrease their overall movement by visual stimulus habituation (Baier & Scott, 2024). Additionally, in comparison to other methods of metabolic activity measurement, such as high-precision respirometry, the alamar assay used here can be affected by other factors, such as the capacity of chemicals to interfere with antioxidant enzymes and molecules (Rajak & Ganguly, 2023). The release of contaminant metabolites that may compete and/or bind with reagents in the assay could also affect the alamar blue results, as this assay works by reducing resazurin to a fluorescent compound resorufin (Munshi et al., 2014). Overall, a chemical causing an increase in oxidative stress would deplete the antioxidant capacity of the fish: this would be observed as a decrease in metabolism in the alamar assay and may not reflect solely the metabolic activity. The alamar blue assay is reflective of the mitochondrial metabolism, while the behavior assay may indicate neurodevelopment toxicity or transient behavior modifications. By the end of the energy expenditure testing, alamar test fish have also been out of chemical exposures for 16 hours, whereas behavioral testing is performed immediately following cessation of the exposures. Thus, this could contribute to differences by itself, and further research should repeat these experiments with the chemical exposures continuing through the various metabolic health testing to elucidate the potential influence of this on the outcomes. A previous study also demonstrated that the swimming activity was affected by the cohort or the exposure round/breeding event (Chackal et al., 2022), which could also have influenced our results. It is also notable that zebrafish behavioral testing continues to improve, with recent protocols for assessing a range of behavioral phenotypes, as well as learning and memory, as well as methods for better delineating mechanisms of effects (Gutsfeld et al., 2024; Leuthold et al., 2024).

There is a very limited number of studies focused on evaluating the effects of organic and inorganic chemicals in mixtures, although they often occur in combination in the environment. Previous studies focused on simple mixtures of two or three chemicals (Di Paola et al., 2021; Kim et al., 2011), or focused on mortality as the main endpoint (Nilén et al., 2022). The study by Kim and collaborators (2011) evaluated the effects of cadmium and PFOS on thyroid and oxidative stress related effects. They observed that a pre-exposure to PFOS increased cadmium toxicity and altered thyroid functions, but the mixture was not more potent than the individual chemical for most of the endpoints. Nilén and collaborators (2022) examined mortality induced by mixtures of increasing complexity using benzo[a]pyrene, PFOS, 3,3’,4,4’,5-pentachlorobiphenyl 126 (PCB-126), and sodium arsenate (As) to evaluate the predictability of the concentration addition and the independent action models. The authors determined that the concentration addition model was more reliable, although the chemicals in their study acted through different modes of action (Nilén et al., 2022). Indeed, the metabolism and detoxification of organic and inorganic chemicals are performed through different pathways, as organic chemicals are detoxified via AHR signaling (Larigot et al., 2018) while the inorganics are metabolized and sequestered by metallothionein (Chan et al., 1989; Chan et al., 2006; Chan, 2023). In our previous work, we evaluated the *in vitro* effects of those chemicals and mixtures on triglyceride accumulation, pre-adipocyte proliferation and receptors bioactivities. We observed a synergistic effects and greater activities from the mixtures compared to the individual components (Bérubé et al., 2023). Although the endpoints measured in both studies differ, the loss of potency in the mixture from this current work is intriguing. The observed difference may be due to the organismal capacity to metabolize and excrete the chemicals or from the chorion protection. Lastly, differences of fish and mammalian metallothionein needs to be evaluated further to compare their capacity to decrease metal toxicity (Capasso et al., 2003).

Previous studies observed other sublethal effects induced by the chemical mixtures (e.g., loss of balance during swimming or lack of swim bladder inflation) that were not induced by the single chemical exposures (Nilén et al., 2022). Our research focused on metabolic-related endpoints, and we did not measure these sublethal effects in this current work. The endpoints chosen in this study may affect the predictability of the mixture compared to the individual components, as the endocrine disrupting effects of these chemicals may not be dose dependent (Hill et al., 2018) and the diverse chemicals examined here have varied pathways of activity, metabolism, and elimination. As noted above, we examined a broad set of both organic and inorganic contaminants as well as several mixtures comprised from these; this limited our ability for more extensive dose response testing of any individual exposure. We instead opted to examine the same four concentrations across all chemicals and mixtures, limiting our ability to assess whole dose responses and complete more comprehensive mixture assessments. Further research can use these broader screening results to focus on specific contaminants and mixtures in a more comprehensive manner to more clearly delineate mixture effects and differences observed here. Lastly, as we did not have a broad enough dose response and/or effect curves to conclusively examine mixture effects with available models, this should instead be viewed as an exploratory extension of our previous *in vitro* mixture assessment of these contaminant combinations, and further studies should examine later life endpoints and should focus in on specific mixtures for more comprehensive evaluations than were possible in this broad study.

## 5. Conclusion

The mixture work performed here is a first step to assessing the endocrine and metabolism disrupting effects from organic and inorganic chemicals in mixtures that represent human exposures to realistic and complex everyday mixtures such as household dust. We observed that mixtures, whether they contained one type of chemical or a combination of organic and inorganic chemicals, did not always reflect the effects of the individual component. Additional work is necessary to fully understand the interactions of the chemicals in the mixture and the effects of those interactions on the toxicity and metabolic health endpoints through more detailed screening of specific mixtures and health effects. Lastly, this work presented the early life metabolic health toxicity representing immediate effects following the exposures. Future work will report the effects of these chemicals and mixtures at later life stages.

## Notes

Funding: Project supported by awards (R00 ES030405 and P30 ES036084) from the National Institute of Environmental Health Sciences (NIEHS), a postdoctoral fellowship from the Natural Sciences and Engineering Research Council of Canada (NSERC; funding reference number 578118) to RB, as well as postdoctoral support for MKL through the Faculty Competition for Postdoctoral Fellows program at Wayne State.

### Competing Interest Statement

The authors have declared no competing interest.

### Summary of Updates

The ORCID number of Dr. Matthew LeFauve was added.

